# Host-adaptation in *Legionellales* is 2.4 Ga, coincident with eukaryogenesis

**DOI:** 10.1101/852004

**Authors:** Eric Hugoson, Tea Ammunét, Lionel Guy

## Abstract

Bacteria adapting to living in a host cell caused the most salient events in the evolution of eukaryotes, namely the seminal fusion with an archaeon ^1^, and the emergence of both the mitochondrion and the chloroplast ^2^. A bacterial clade that may hold the key to understanding these events is the deep-branching gammaproteobacterial order *Legionellales* – containing among others *Coxiella* and *Legionella* – of which all known members grow inside eukaryotic cells ^3^. Here, by analyzing 35 novel *Legionellales* genomes mainly acquired through metagenomics, we show that this group is much more diverse than previously thought, and that key host-adaptation events took place very early in its evolution. Crucial virulence factors like the Type IVB secretion (Dot/Icm) system and two shared effector proteins were gained in the last *Legionellales* common ancestor (LLCA), while many metabolic gene families were lost in LLCA and its immediate descendants. We estimate that LLCA lived circa 2.4 Ga ago, predating the last eukaryotic common ancestor (LECA) by at least 0.5 Ga ^4^. These elements strongly indicate that host-adaptation arose only once in *Legionellales*, and that these bacteria were using advanced molecular machinery to exploit and manipulate host cells very early in eukaryogenesis.

## Introduction

The recent discovery of Asgard archaea and their placement on the tree of life ^5,6^ sheds light on early eukaryogenesis, confirming that eukaryotes arose from a fusion between an archaeon and a bacterium. However, many questions pertaining to eukaryogenesis remain unanswered. In particular, the timing and the nature of the fusion is vigorously debated ^1,7–9^. Mito-late scenarios ^8,10,11^ posit that endosymbiosis of the mitochondrion occurred in a eukaryote that was capable of phagocytosis, while mito-early scenarios ^12^ posit that mitochondrial endosymbiosis triggered eukaryogenesis. In many scenarios, phagocytosis is considered a prerequisite for mitochondrion endosymbiosis and therefore a key component needed for eukaryogenesis ^7^. It is not certain when eukaryotes gained the ability to phagocytose bacteria, but it was most certainly prior to the last eukaryotic common ancestor (LECA) ^1,7^.

The rise of bacteria-phagocytosing eukaryotes created a new ecological opportunity for bacteria, as the eukaryotic cytoplasm is a rich environment. Many bacteria have adapted to exploit this niche, by resisting digestion once phagocytosed by their predators. A host-adapted lifestyle has evolved many times in many different taxonomic groups ^13^, but there are few large groups comprised solely of host-adapted members; such groups include *Legionellales*, *Chlamydiales*, *Rickettsiales* and *Mycobacteriaceae*. *Legionellales* is a diverse group of *Gammaproteobacteria* ^3,14^ that lives only intracellularly and include the well-studied accidental human pathogens *Coxiella burnetii* and *Legionella* spp. *Legionellales* species vary greatly in lifestyle ^3^, ranging from facultatively intracellular (like *Legionella* spp.) to obligatorily intracellular (like the vertically inherited *Coxiella* symbionts of ticks ^15,16^). Because of their rarity and fastidious nature, these host-adapted bacteria are underrepresented in genomic databases; individual isolates from only 7 genera have been sequenced out of an estimated > 450 genera ^14^.

Hallmarks of adaptation to intracellular space include smaller population sizes, genome reduction and degradation, and pseudogenization ^13^. In addition, the infection process linked to an intracellular lifestyle requires a number of specialized functions. For instance, the Type IVB secretion system (T4BSS) plays a key role in the interaction of *Coxiella* ^17^ and *Legionella* ^18,19^ with their hosts, injecting a wide diversity of protein effectors into the host cell. These effectors alter the host behavior, preventing the bacterium from being digested and aiding exploitation of host resources, among others. Some *Legionella* species encode as many as 18 000 types of effectors, but only eight of these are conserved throughout the genus ^20,21^.

To better understand the evolutionary history of *Legionellales* and its relationships with their early hosts, we gathered 35 genomic sequences from novel *Legionellales* through whole-genome sequencing, binning of metagenome-assembled genomes (MAGs), and database mining.

## Results and Discussion

### Phylogenetic relationships and diversity among *Legionellales*

Bayesian and maximum-likelihood phylogenies confirm that *Legionellaceae* and *Coxiellaceae* are monophyletic sister clades (hereafter jointly referred as *Legionellales* sensu stricto) are monophyletic, sister clades, and diverged rapidly from each other after the last *Legionellales* common ancestor (LLCA) (**Fig. 1**). The phylogeny reveals extensive, previously unknown diversity among *Legionellales* (**Fig. 1**). *Coxiellaceae* comprises two large clades, one including the known genera *Coxiella*, *Rickettsiella* and *Diplorickettsia*, and the other including the amoebal pathogen *Aquicella*^22^ and environmental MAGs. Notably, the two MAGs in *Legionellales* that are resolved as outside *Legionella* have small genomes (2.2 and 1.9 Mb, respectively, see **Supp. Figure 3**), whereas genomes within *Legionella* (except for *Ca.* L. polyplacis, an obligate louse endosymbiont ^23^) range from 2.3 to 4.9 Mb ^21^. This pattern suggests that the last common ancestor of *Legionellaceae* had a smaller genome than extant *Legionella* species, although it cannot be completely excluded that genome reduction occurred independently in these two MAGs.

**Figure 1.**
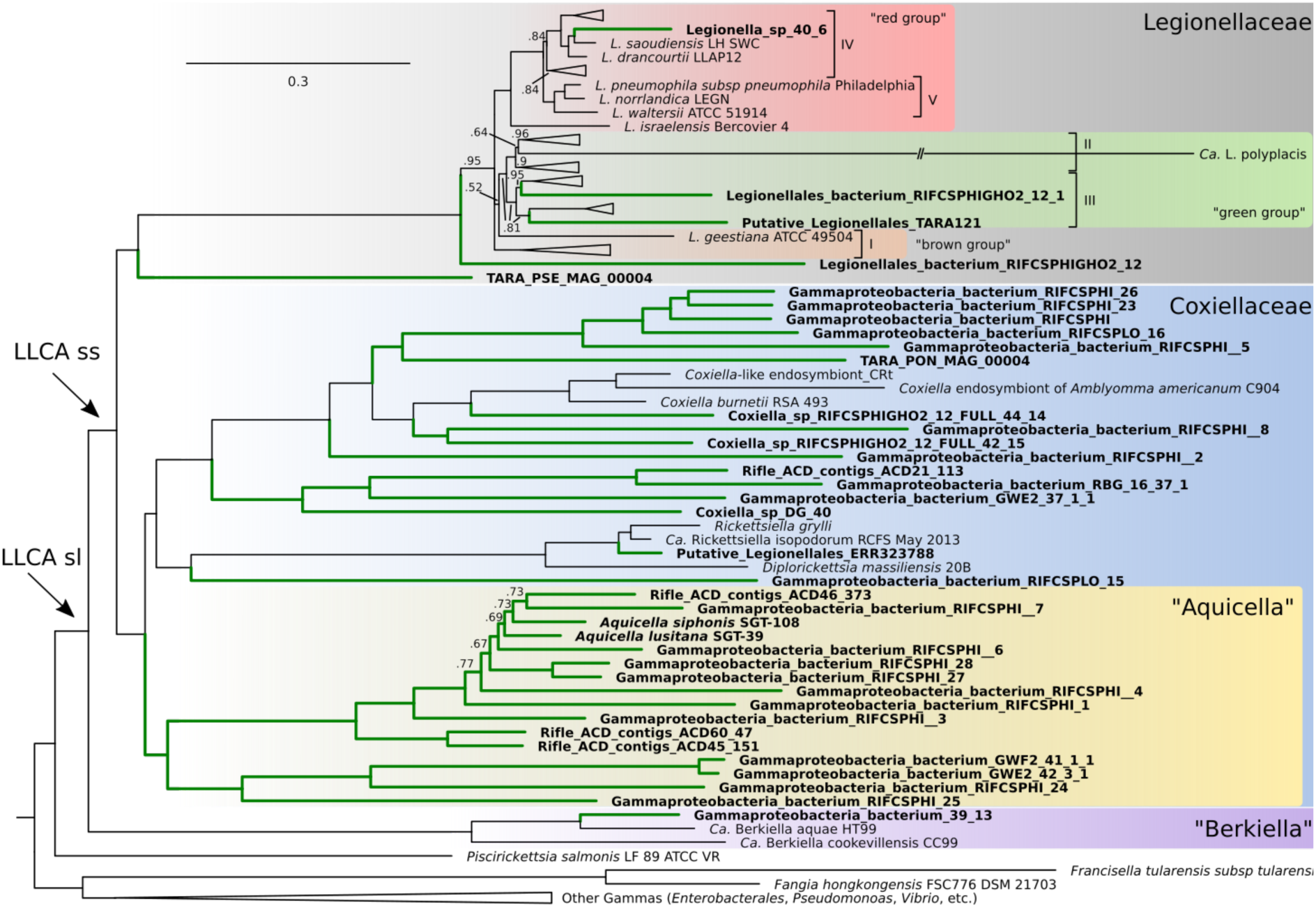
Bayesian phylogeny of *Legionellales*. Tree inferred with PhyloBayes using a GTR+CAT model and a concatenated amino acid alignment of 109 single-copy orthologs (Bact109) comprising 93 *Legionellales* genomes, 16 genomes from other *Gammaproteobacteria*, and rooted with an outgroup of 4 genomes from non-*Gammaproteobacteria Proteobacteria* (dataset Legio93). Numbers on branches show posterior probabilities (pp). Branches without numbers indicate pp = 1. Terminal nodes in bold indicate taxa recovered in this study either by database mining or sequencing. Green branches indicate clades where no genomic data was available prior to this study. The scale shows the average number of substitutions per site. The colored groups and Roman numerals in *Legionellaceae* correspond to the groupings in ^20^ and ^21^, respectively. The full tree is available in **Supp. Figure 1**.

Interestingly, two MAGs recovered from the TARA Ocean sampling campaign are included in the *Legionellaceae*, one as a sister clade to the *Legionella* genus, and one clustering within the “green group”, having *L. birminghamensis* and *L. quinlivanii* as closest relatives (**Fig. 1**). The discovery of two *Legionella* genomes in a marine environment supports rising evidence that *Legionella* and other genera of the *Legionellales* can colonize saline waters ^14,77^. Strikingly, both MAGs branching as sister groups to the *Legionella* genus have small genomes (2.2 and 1.9 Mb, respectively), whereas genomes in the *Legionella* genus (except for *Ca.* L. polyplacis, which is an obligate endosymbiont of a louse ^23^) range from 2.3 to 4.9 Mb ^21^. This argues in favor of a *Legionellaceae* last common ancestor that had a smaller genome than the extant *Legionella* species. Incidentally, both MAGs contained inside the *Legionella* clade III ^21^ also display smaller genomes than their closest relatives: 1.9 and 2.3 Mb, whereas clade III *Legionella* spp. range from 3.1 to 3.9 Mb. The same is also true, albeit to a lesser extent, with the MAG branching in clade IV (Legionella_sp_40_6: 3.1 Mb; rest of the clade: 3.4-4.9 Mb). The reduced genome size in these MAGs, associated with the longer branches leading to them, might indicate shifts in lifestyles, either towards streamlining as in the *Pelagibacterales* ^78^ or towards an increased dependency on their host ^13^, similarly to *Ca.* Legionella polyplacis ^23^.

Two other groups, *Berkiella* spp. and *Piscirickettsia* spp. could not be placed with very high confidence in the tree (**Supp. Table 1**). However, Bayesian phylogenies (**Supp. Figure 1 and 2**) inferred with the CAT model consistently placed *Berkiella* as sister clade to *Legionellales* sensu stricto (ss), so *Berkiella* + *Legionellales* is hereafter referred to as *Legionellales* sensu lato (sl). The placement of *Piscirickettsia* is not as consistent, and further studies are needed to establish whether it is more closely related to *Legionellales* or the *Francisella*/*Fangia* group.

### Evolution of genome content in *Legionellales*

To better understand the evolution of host-adaptation in the order, we reconstructed gene flow within *Legionellales*. The phylogenetic birth-and-death model implemented in Count ^24^ was used on the same set of genomes (Legio93) as the tree in **Fig. 1**. The reconstructed gene flow reveals 553 gene family losses from the last free-living ancestor to LLCA sl (**Fig. 2, Supp. Figure 3, Supp. Table 2**). These 553 losses can be compared to 781 losses on the branch leading to *Fangia*/*Francisella*, another intracellular group. A subsequent large gain in gene families in the last common ancestors of *Legionellaceae* and *Berkiella* spp. (588 and 556, respectively) but not in the ancestors of *Coxiellaceae* (*Coxiella/Aquicella*) suggests that LLCA had a genome size comparable to that of the latter group (average: 1.8 Mb), compatible with genome sizes of gammaproteobacterial intracellular bacteria ^13^.

A crucial feature for the success of many host-adapted bacteria is the ability to transfer proteins into the host’s cytoplasm, typically through secretion systems. Among the proteobacterial VirB4-family of ssDNA conjugation systems (Type IV Secretion Systems, T4SS), one called T4BSS (also called MPF_I_ or I-type T4SS) probably arose early in the group ^25^. It is present in almost all extant *Legionellales*, with the exception of the extremely reduced endosymbionts of the group (**Supp. Figure 3**), where it is either missing completely or pseudogenized. It is partly missing in several of the small, novel *Coxiellaceae* MAGs, possibly indicating increased host dependence. However, it is difficult to assess with certainty whether genes absent in MAGs represent true losses or are the result of incomplete binning of metagenomic contigs. T4BSS in *Legionellales* appears largely collinear (**Fig. 3, Supp. Figure 4**). Ancestral reconstruction of the order revealed that T4BSS, as it is found in extant *Legionellales*, originated after the last free-living ancestor but before the LLCA, with several proteins added over time (**Fig. 2**). IcmW and IcmS (two small acidic cytoplasmic proteins, part of the coupling protein subcomplex ^79,80^), IcmC/DotE, IcmD/DotP and IcmV (three small integral inner membrane proteins, of uncharacterized functions) were gained in the LLCA sl. IcmQ, a cytoplasmic protein and IcmN/DotK, an outer membrane lipoprotein ^81^, were acquired in the LLCA sl stricto, after the divergence with *Berkiella*. The limited amount of characterized functions for these proteins prevents us to infer exactly how their gain allowed the LLCA to infect eukaryotic cells. However, except for IcmN/DotK, all these proteins are located either in the inner membrane or in the cytoplasm, suggesting that the specific role of the *Legionellales*-gained proteins is related to the coupling protein complex rather than in the core complex.

**Figure 2.**
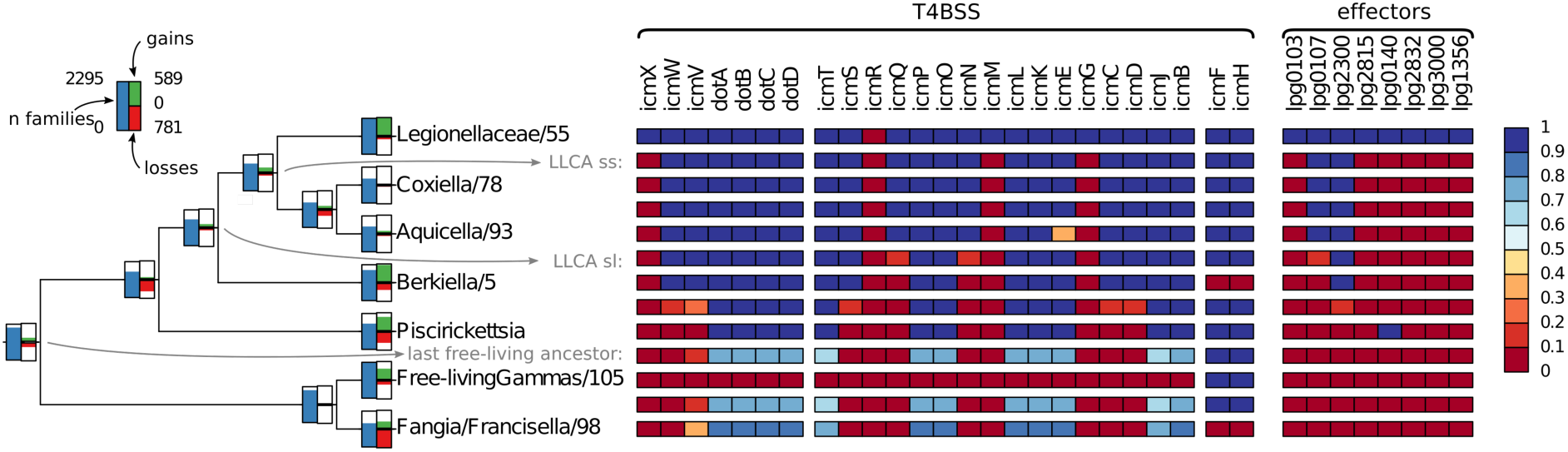
Overview of ancestral reconstruction of *Legionellales* genomes, with the T4BSS system and the *Legionella*-conserved effectors detailed. The ancestral reconstruction is based on the Bayesian tree in Fig. 1. At the left of each node, barplots depict the number of gene families (blue, ranging from 0 to 2295) inferred at that node, as well as the number of gene family gains (green, from 0 to 589) and losses (red, from 0 to 781) on the branches leading to nodes. On the right panel, squares represent genes in the T4BSS (separated according to the three operons present in R64 ^19^) and the eight effector genes conserved in *Legionella* ^21^. Rows in-between terminal nodes correspond to the last common ancestor of the two nearest terminal nodes, e.g. the row between *Aquicella* and *Berkiella* corresponds the LLCA ss. The color of the squares represents the posterior probability that each protein was present in the ancestor of each group. A complete tree is shown in **Supp. Figure 3** and the underlying numbers are available in **Supp. Tables 2 and 3**.

**Figure 3.**
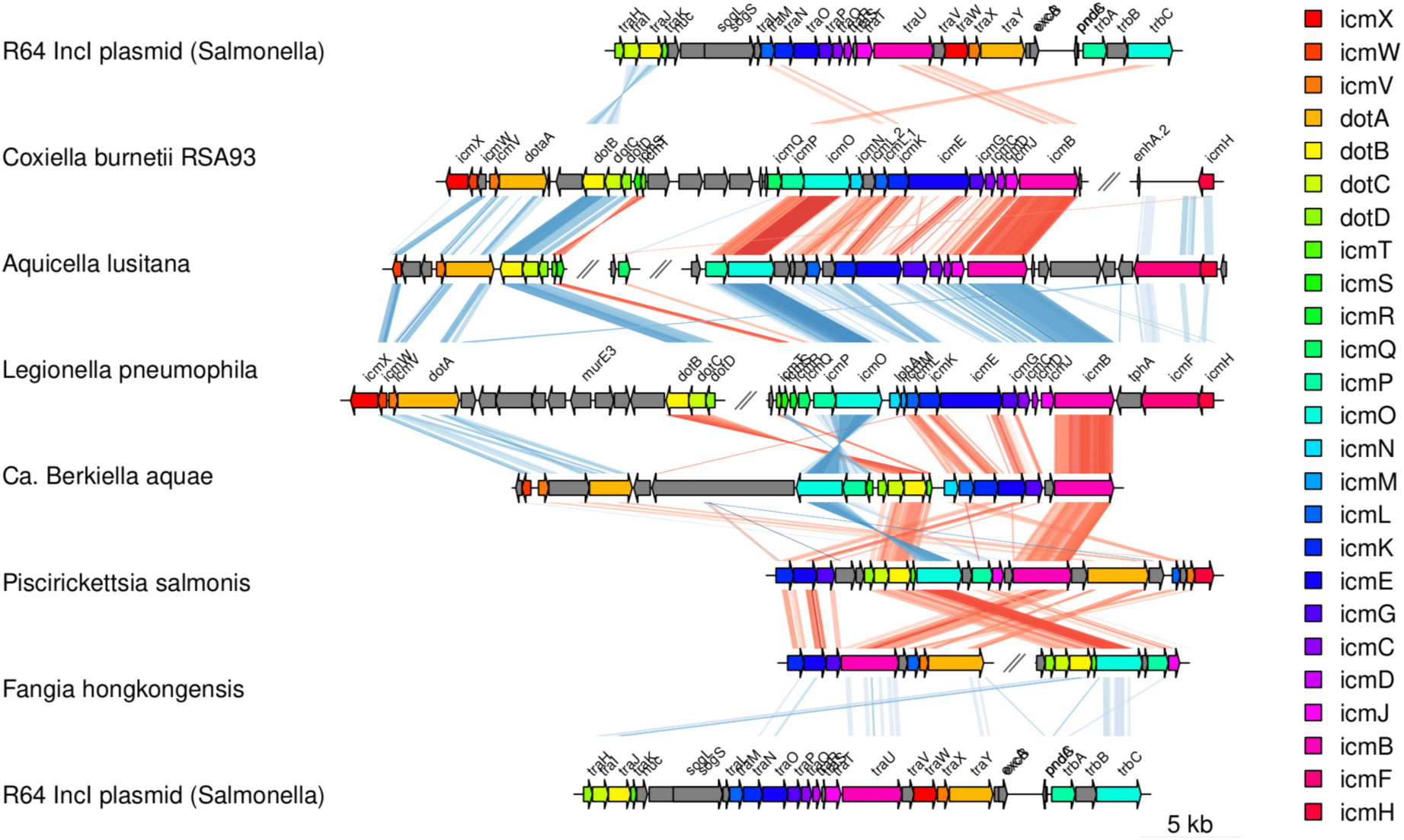
Collinearity of T4BSS in *Legionellales*. Genes are colored as they appear in *Legionella pneumophila* Philadelphia. Lines connecting rows show similarities between the sequences, as identified by tblastx. Red lines indicate a direct hit, blue lines a complementary one. Darker shades indicate higher bit scores of the tblastx hit.

A homologous T4BSS system is present in *Fangia hongkongensis*, but mostly absent in the free-living *Gammaproteobacteria* and in *Francisella*, a sister clade to *Fangia*. It is likely that the T4BSS in *Fangia* has been acquired by horizontal gene transfer, possibly from *Piscirickettsia* (**Supp. Figure 5** and **6**).

Contrarily to T4BSS itself, which is highly conserved throughout the order, effectors are much more versatile. Of the eight effectors conserved in the *Legionella* genus ^21^, only two are found outside of the *Legionellaceae* family (**Fig. 2**): RavC (lpg0107) is found in most *Legionellales* ss, while LegA3/AnkH/AnkW (lpg2300) is found in *Legionellales* ss and *Berkiella*. MavN (lpg2815), which is conserved in *Legionella* spp. and was found in *Rickettsiella* ^20^, is found only in a few *Coxiellaceae* species (**Fig. 2**; **Supp. Figure 3**), and is unlikely to have been acquired in the LLCA. Although the function of these effectors is not precisely known, LegA3/AnkH contains ankyrin repeats, and has been suggested to play a role in process involved in modulation of phagosome biogenesis ^26^.

### Dating the rise of the *Legionellales*

Absolute dating of bacterial trees is often inconclusive, because very few reliable biomarkers can be unambiguously attributed to a bacterial clade and used as calibration points ^27^. However, one such biomarker is okenone, a pigment whose degradation product okenane has been found in the 1.64-Ga-old Barney Creek Formation ^28^, and is exclusively produced by a subset of *Chromatiaceae*. *Chromatiaceae* (part of the purple sulfur bacteria) and *Legionellales* are phylogenetically close to each other within *Gammaproteobacteria* ^29^. We reasoned that the last common ancestor of all okenone-producing *Chromatiaceae* had to be at least as old as the earliest known trace of okenone, i.e. ≥1.64 Ga (**Fig. 4**). Using this constraint and a relaxed molecular clock model, we were able to calibrate a Bayesian tree based on a dataset including genomes from 105 *Gammaproteobacteria* (of which 22 belong *Legionellales* and 19 to *Chromatiaceae*) and 5 outgroups (Gamma105 dataset). We estimate that LLCA existed ca. 2.43 Ga ago (**Fig. 4**), with a 95% high posterior density ranging from 2.70 to 2.26 Ga (see **Supp. Table 4**).

**Figure 4.**
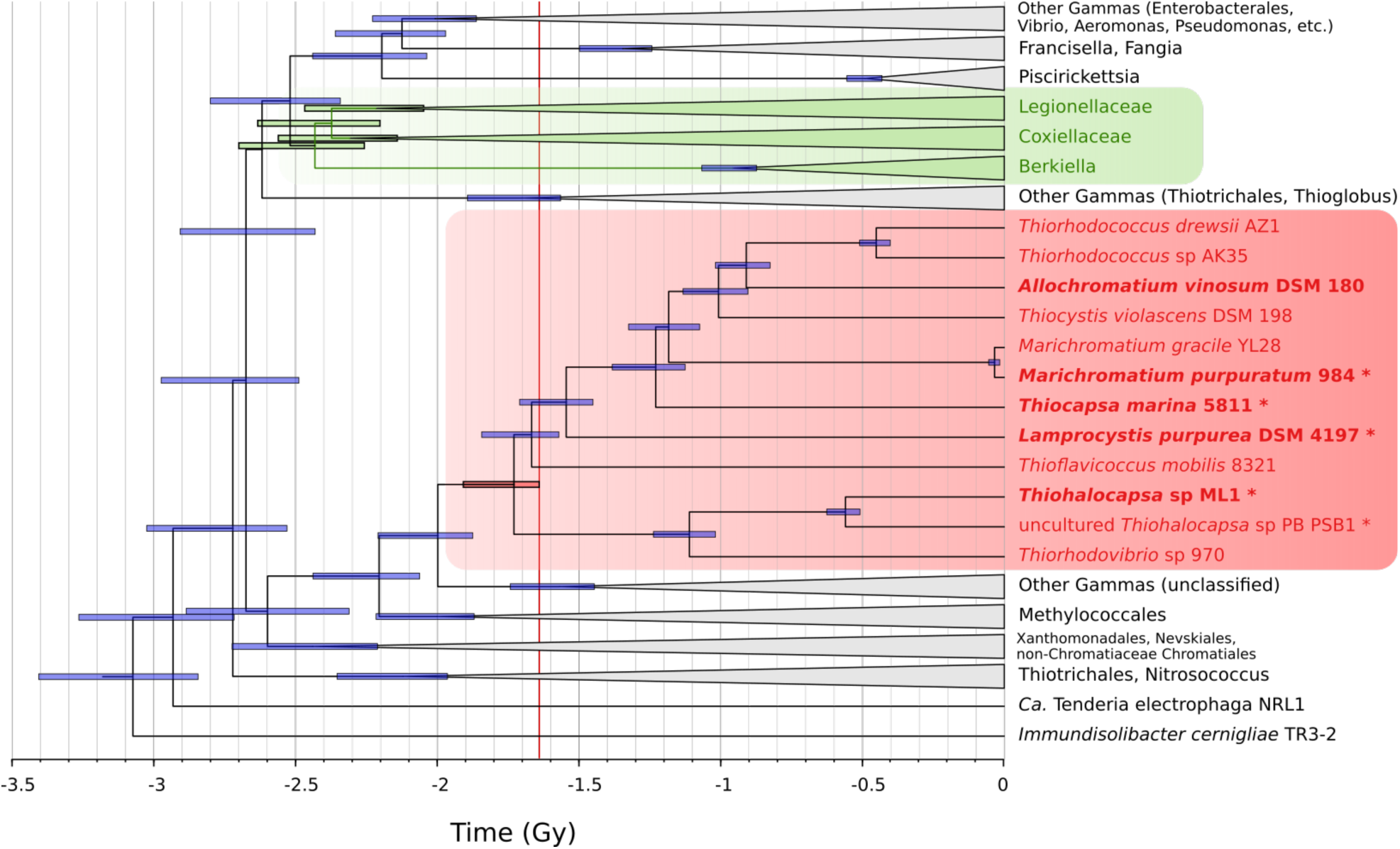
Bayesian estimation of the appearance of the LLCA. *Chromatiaceae* are highlighted in red, with okenone-producing species in bold. *Legionellales* are highlighted in green. The red vertical line is drawn at 1.64 Ga, the age of the Barney Creek Formation which contains okenane, which gives the lower estimate for the divergence of *Chromatiaceae*. Horizontal bars at interior nodes represent the highest posterior density interval (95%) as determined by BEAST. The red horizontal bar indicates the last common ancestor of all okenone-producing *Chromatiaceae*, the green bars indicate ancestors of *Legionellales*, with, from top to bottom, the last common ancestor of *Legionellaceae*, *Legionellales* ss*, Coxiellaceae*, and *Legionellales* sl (LLCA). The underlying tree is shown in **Supp. Figure 2**.

We took several steps to mitigate confounding factors in estimating the age of LLCA. Firstly, we carefully selected genomes included in the Gamma105 dataset: (i) we included a large outgroup, both in the *Gammaproteobacteria* and outside; (ii) whenever possible, we included genomes that were breaking long branches; (iii) we removed fast-evolving genomes, but kept those breaking long branches.

Secondly, we tested several clock models: strict, relaxed with uncorrelated log-normal distribution, and relaxed with random local distribution. The latter was not computationally tractable. The four chains run for the strict clock model (10 million generations each) converged but the strict clock assumption appeared invalid when examining the variation of mutation rates obtained from the relaxed clock model. The results of the strict clock model were therefore not used. For the relaxed clock model (uncorrelated log-normal), five chains were run, one for 55.7 million generations, and four for 10 million generations each. Most of the parameters achieved an ESS score > 100, a good indication that the chains had efficiently sampled the parameter space. The mutation rate and the mean of the ucld parameter did not reach high ESS scores but were negatively correlated with each other. Despite not fully converging, the five trees from the five chains yielded similar results. In particular, the estimation of the age of LLCA was consistent between the chains (see **Supp. Table 4**), which strongly indicates that the chains correctly explored the parameter space. To estimate how the chosen priors affected the runs, the parameters obtained from the actual run (57 million generations) were compared with those from a run with 10 million generations drawing only from the priors. All parameters displayed very different distributions, suggesting that the choice of the priors had very little effect on the obtained values.

## Conclusion

The common ability of *Legionellales* bacteria to invade eukaryotic cells combined with the presence and high conservation of the crucial host-adaptation genes, namely a complete T4BSS and two effectors, strongly suggest that the last common ancestor of *Legionellales* was already infecting other cells. The hypothesis that LLCA used these mechanisms to infect or interact with other prokaryotes cannot be completely ruled out, but seems extremely unlikely for several reasons. For example, if LLCA was only infecting prokaryotes, then the ability to infect eukaryotes would have to have evolved independently several times within *Legionellales*, in all free-living descending clades, with no exception (all experimentally investigated *Legionellales* are able to live inside the cytoplasm of eukaryotic cells). Furthermore, each of these independent host-adaptation events would have to have involved the same host-adaptation genes (all experimentally investigated *Legionellales* use the same T4BSS to interact with their hosts).

It is also likely that LLCA, like extant *Legionellales*, depended on phagocytosis (or its prototypic version), a mechanism exclusively found in eukaryotes. This implies that LLCA appeared after the division of Archaea and the first eukaryotic common ancestor (FECA), thus placing rise of phagocytosis and FECA at 2.26 Ga at the latest. Given a consensual estimate of the last eukaryotic common ancestor’s (LECA) age of 1.0–1.6 Ga ^4^, the time for eukaryogenesis, defined as the time elapsed between FECA and LECA ^1^, would be at least 660 Ma, but could be >1.2 Ga.

The existence of a phagocytosing eukaryote at 2.26 Ga has implications for hypotheses of eukaryogenesis. Four of the most prominent of these (reviewed e.g. in ^9^) make different assumptions about the timing of the mitochondrial endosymbiosis. In the hydrogen hypothesis ^12^, the mitochondria arrived early (‘mito-early’), with the mitochondrial endosymbiosis event itself triggering eukaryogenesis. In the phagocytosing archaeon model (PhAT) ^30^, the syntrophy hypothesis ^10^, and the serial endosymbiosis model ^8^, the mitochondrion arrive late (‘mito-late’). Specifically, in the PhAT model, phagocytosis machinery is a pre-requisite for the fusion of the mitochondrial progenitor with an Asgard archaeon. The very early emergence of phagocytosis proposed in this contribution supports a mito-late scenario.

In summary, we show here that the last *Legionellales* common ancestor already possessed a T4BSS and two effectors, and existed over 2.26 Ga ago. We propose that LLCA, upon being phagocytosed by eukaryotic cells, already had the ability to resist digestion, owing to its host-adaptation genes. Thus, phagocytosis is at least 2.26 Ga old, earlier than previously thought. This hypothesis is consistent with a scenario in which some early eukaryotes developed phagocytic properties and fed on prokaryotes. Some of these, among them LLCA, rapidly acquired the abilities to resist host digestion and exploit the novel, rich ecological niche that is the eukaryotic cytoplasm.

## Material and Methods

### Bacterial strains

Two species of *Aquicella* (*A. lusitana*, DSM 16500 and *A. siphonis*, DSM 17428) ^22^ were ordered from the DSMZ strain culture and grown at 37°C for 5 days on Buffered Charcoal Yeast Extract (BCYE) agar plates with *Legionella* BCYE supplement (OXOID).

### DNA Extraction and sequencing, assembly and annotation

DNA was extracted from pure culture isolates (*Aquicella* spp.) using the Genomic Tip 100 DNA extraction kit (QIAGEN), following the manufacturer’s instructions. DNA extracted from pure cultures was sequenced by PacBio, using one SMRT cell for each isolate. Reads were assembled using the HGAP.3 pipeline, resulting in one unitig per replicon. The assembly was confirmed by performing cumulative GC skews on each replicon ^31^. All replicons had repetitive sequences at the beginning and the end of the sequences, suggesting that they were circular. One unitig in *A. lusitana* had multiple repeats of the same sequences and was edited manually. All chromosomes were rotated to have the origin of replication, as determined by cumulative GC skews, at the beginning of the sequence. The final sequences were confirmed by mapping ∼500k shorter reads (2×300 bp, obtained from an Illumina MiSeq run) to the sequences obtained from PacBio. No SNP could be found in any of the replicons. All sequencing, as well as quality control and assembly of the PacBio reads, was performed at NGI, SciLifeLab, Uppsala and Stockholm, Sweden.

The sequences were then annotated with prokka 1.12-beta ^32^, using prodigal 2.6.3 ^33^ to call protein-coding genes, Aragorn 1.2 ^34^ to predict tRNAs and barrnap 0.7 ^35^ to predict rRNAs. Default functional annotation was performed by prokka. Both genomes are submitted at the European Nucleotide Archive (ENA) under study number PRJEB29684.

### Protein marker sets

A set of 139 PFAM protein domains was used as a starting point both to perform quality control of MAGs and for phylogenomic reconstructions. The original set was used by Rinke et al (Supplementary Table 13 in ^36^) to estimate the completeness of their MAGs. We used the same set (referred to as Bact139) to estimate MAG completeness. A large subset of the 139 protein domains are very frequently located in the same protein in a vast majority of Proteobacteria. This was evidenced by investigating 1083 genomes, using phyloSkeleton 1.1 ^37^ to select one representative per proteobacterial genus and to identify proteins containing the Bact139 domain set. Of these domains which often co-localized in the same protein, only the most widespread one was retained. In total, only 109 domains were used to identify proteins suitable for phylogenomics analysis. This set is referred to as Bact109 and is available in phyloSkeleton 1.1 ^37^.

### Metagenome assembly

The 86 metagenomes with the highest fraction of reads belonging to *Legionellales* ^14^ were selected (**Supp. Table 5**) and downloaded from MGnify ^38^. Metagenomes sequenced only with 454 (Roche), or that were too complex to be assembled on the computational cluster at our disposition (512 Gb RAM) were not included. Raw reads were downloaded from the European Nucleotide Archive (ENA) and trimmed using Trim Galore (v0.4.2) ^39^ to remove standard adapter sequences. Additionally, reads were trimmed using Trimmomatic (v0.36) ^40^ with a sliding window (4:15) to only discard low-quality bases without losing whole reads. Trimmed reads were then assembled using SPAdes (St. Petersburg genome assembler) (v3.7.1) ^41^, using the --meta option and kmer sizes 21, 33, and 55 for metagenome sizes that did not require running on a HPCC (High Performance Computer Cluster). More complex metagenomes requiring a HPCC were assembled on UPPMAX with a single kmer, 31. Contigs smaller than 500 bp were then discarded.

### Identification of MAGs belonging to *Legionellales*

To identify MAGs belonging to *Legionellales*, the ggkBase (https://ggkbase.berkeley.edu/) and NCBI Genome databases were screened for MAGs attributed to Gammaproteobacteria. MAGs obtained through metagenomic binning from a previous published study ^42^ and from a prepublication ^43^ that branched close to *Legionella pneumophila* were also included.

The selected MAGs and the assembled metagenomes were screened using the same ribosomal protein (r-protein) pipeline as described in ^6^, hereafter referred to as RP15. Briefly, this pipeline uses first PSI-BLAST 2.9.0+ to search the given metagenomes and MAGs for 15 ribosomal proteins, which are universally located in a single chromosomal locus. Contigs containing at least 8 of the 15 r-proteins are then retained, and referred to as r-contigs.

A backbone of known organisms (with complete genomes and established taxonomies) was used to help identifying to which known clades the r-contigs belong. The package phyloSkeleton 1.1 ^37^ was used to select one representative genome in each family within Gammaproteobacteria, one in each order of the Beta- and Alphaproteobacteria, as well as *Mariprofundus* and *Acidithiobacillus*. phyloSkeleton also identified orthologs of 15 ribosomal proteins in the gathered genomes.

The RP15 orthologs gathered from both the metagenomic sources and the backbone were aligned with MAFFT L-INS-I v7.273 ^44^, and the alignments concatenated through the RP15 pipeline. The resulting concatenated alignment was used to infer a rough phylogenetic tree using FastTree 2.1.8 ^45^, using the WAG model. R-contigs that were clearly unrelated to *Legionellales* were removed from the dataset, while the remaining proteins were realigned and concatenated, to generate a new tree. This process was iterated until all remaining MAGs and r-contigs were deemed belonging to *Legionellales*. A final phylogeny was inferred with RAxML 8.2.8 ^46^ using the PROTGAMMA model which allowed automatic selection of best substitution matrix to infer a more robust phylogeny.

### Metagenomic binning and quality control

Metagenomes containing r-contigs belonging to *Legionellales* were then binned with ESOM 1.1 ^47,48^. An ESOM map was generated from contigs over 2 kb with a 10 kb sliding window, and visualized using the ESOM analyzer, highlighting the location of the r-contig and selecting the corresponding bin. The completeness and redundancy of all MAGs were then controlled using miComplete 0.1 ^49^

Some MAGs were very similar to each other, due to different versions of the same bin being present in databases. The most complete MAG was then selected for each set, provided they were not more redundant than another genome within the same set. If equal in completeness the NCBI-submitted MAG was preferred.

### Phylogenomics

Two sets of genomes were used for phylogenomics in this contribution. Both sets consist of part or all of the novel *Legionellales* MAGs and genomes, supplemented with varying subsets of outgroups. The initial selection of the representative organisms and identification of markers was done with phyloSkeleton 1.1 ^37^, initially choosing one representative per class in the *Betaproteobacteria*, the *Zetaproteobacteria* and the *Acidithiobacillia*; one per genus in the Gammaproteobacteria, except in the *Piscirickettsiaceae* and the *Francisellaceae* (one per species). That initial selection of genomes was then trimmed down to reduce the computational burden of tree inference.

For both sets, protein markers from the Bact109 set were identified with phyloSkeleton 1.1 ^37^. Each marker was aligned separately with mafft-linsi v7.273 ^44^. The resulting alignments were trimmed with BMGE v1.12 ^50^, using the BLOSUM30 matrix and the stationary-based trimming algorithm. The alignments were then concatenated. From these alignments, maximum-likelihood trees were reconstructed with FastTreeMP v2.1.10 ^45^ using the WAG substitution matrix. Upon visual inspection of the resulting trees, the genome selection was reduced to remove closely related outgroups, and the phylogenomic procedure repeated. Once the genome dataset was final, a maximum-likelihood tree was inferred with IQ-TREE v1.6.5 ^51^, using the LG ^52^ substitution matrix, empirical codon frequencies, four gamma categories, the C60 mixture model ^53^ and PMSF approximation ^54^; 1000 ultrafast bootstraps were drawn ^55^.

The first set of genomes, called Gamma105, was used to correctly place *Legionellales* in *Gammaproteobacteria*, in particular with respect to *Chromatiaceae*. It encompasses 110 genomes, of which 105 are *Gammaproteobacteria*, one is a *Zetaproteobacterium*, 3 are *Betaproteobacteria*, and one is an *Acidithiobacillia*. The selected *Gammaproteobacteria* include representatives from *Legionellales* (22), *Chromatiaceae* (19), *Francisellaceae* (17) and *Piscirickettsiaceae* (4) (**Supp. Table 6**).

The second set, called Legio93, is focused on *Legionellales* itself and comprises 113 genomes, of which 93 belong to *Legionellales* and the rest to other *Gammaproteobacteria* (16 genomes), *Betaproteobacteria* (2 genomes), *Acidithobacillia* (1 genome) and *Zetaproteobacteria* (1 genome) (**Supp. Table 7**).

For both sets, single-gene maximum-likelihood trees were inferred for each marker. The BMGE-trimmed alignments were used to infer a tree using IQ-TREE 1.6.5, using the automatic model finder ^56^, limiting the matrices to be tested to the LG matrix and the C10 and C20 mixture models. Single-gene trees were visually inspected for the presence of very long branches resulting from distant paralogs being chosen by phyloSkeleton.

For both sets, a Bayesian phylogeny was inferred with phylobayes MPI 1.5a ^57^ from the concatenated BMGE-trimmed alignments, using a CAT+GTR model. Four parallel chains were run for 9500 generations (Gamma105 set) and 3500 generations (Legio93 set), respectively. The chains did not converge in either set. For Legio93, all four chains yielded the same topology for all deep nodes in and around *Legionellales*. For Gamma105, 3 out of 4 chains yielded the same overall topology, while the last one had the *Piscirickettsia* and the *Berkiella* clades inverted (**Supp. Table 1**). The majority-rule tree obtained from the Bayesian trees for Legio93 was subsequently used for the ancestral reconstruction (see below), while the one for Gamma105 was used as input tree to estimate the time of divergence of the LLCA (see below).

### Identification of okenone-producing *Chromatiaceae*

To relate the earliest documented trace of okenone ^28,58^ to a specific ancestor, we took two approaches. First, we compared a list of okenone-producing *Chromatiaceae* ^59^ with the genomes available in Genbank and at the JGI Genome Portal ^60^. We found five okenone-producing *Chromatiaceae* genomes. Second, we used homologs of the proteins encoded by the genes *crtU* and *crtY*, which are considered essential to synthesize okenone ^61^, to screen the nr database. We used the proteins from *Thiodictyon syntrophicum* str. Cad16 (accession number: CrtU, AEO72326.1; CrtY, AEO72327.1) as queries in a PSI-BLAST 2.9.0+ search ^62^ restricted to the order *Chromatiales*, keeping only proteins with at least 50% similarity. Gathered homologs (n=7 in both cases) were aligned with MAFFT L-INS-I v7.273 ^44^ and the alignment was pressed into a hidden Markov model (HMM) with HMMer 3.1b2 ^63^. Using phyloSkeleton ^37^, we collected all *Chromatiales* genomes from NCBI (n=130) and queried these, together with the Gamma105 set with both HMMs. A significant similarity (E-value < 10^-10^) for CrtU and CrtY was found in 42 and 25 genomes, respectively. A hit for both proteins was found in 8 genomes only, of which 6 belong to the *Chromatiaceae*. Of these 6, 4 were already identified by the first method listed above; one was a MAG that was too incomplete to include in further phylogenomics analysis; the last one was another MAG that was included in later analysis. The other two genome that had proteins similar to CrtU and CrtY are *Kushneria aurantia* DSM 21353, a member of the *Halomonadaceae* and *Immundisolibacter cernigliae* TR3-2, which branches very close to the root of the *Gammaproteobacteria*: these two might represent cases of horizontal gene transfer.

### Time-constrained tree

To estimate the time of divergence of LLCA, we used BEAST 2 ^64^, using a strict and a relaxed clock model (uncorrelated log-normal) ^65^. To reduce computational load, we used a fixed input tree (Bayesian, Gamma105; see above). The following parameters were used: the concatenated alignment (104 proteins) was used as a single partition; for site model, a LG matrix with 4 gamma categories and 10% invariant was used, estimating all parameters; for the relaxed clock model, an uncorrelated log-normal model was used, estimating the clock rate. The following prior distributions were used: shape of the gamma distributions for the site model: exponential; mutation rate: 1/x; proportion of invariants: uniform; ucld mean: 1/X, offset: 0, ucld std: gamma. To calibrate the clock, it was assumed that the last common ancestor of all okenone-producing *Chromatiaceae* was at least as old as the rocks where the earliest traces of okenane (a degradation product of okenone) was found, 1.640 ± 0.03 Ga old ^28,58,66^. A prior was thus set for that the divergence time of the monophyletic group comprising these organisms: the divergence time was drawn from an exponential distribution of mean 0.1 Ga with an offset of 1.64 Ga. Although it cannot be completely excluded, no evidence suggests that other organisms had produced okenone earlier than the last common ancestor of *Chromatiaceae*: the only mentions of okenone in the scientific literature are linked to the presence of *Chromatiaceae*.

The four chains run for the strict clock model (10 million generations each) converged but the strict clock assumption seemed unrealistic when examining the variation of mutation rates obtained from the relaxed clock model. For the relaxed clock model, five chains were run, one for 55.7 million generations, and four for 10 million generations each. Traces were examined with Tracer 1.6.0 ^67^. Most of the parameters achieved an ESS score > 100. The mutation rate and the mean of the ucld parameter did not reach very high ESS scores but were negatively correlated with each other. Trees were extracted with TreeAnnotator 2.4.7 (part of BEAST), deciding the burn-in fraction after visual examination of the traces (see **Supp. Table 4**). Despite not fully converging, the five trees from the five chains yielded similar results.

To estimate how the chosen priors affected the runs, a run with 10 million generations was performed, drawing from the priors. Parameters were compared in Tracer 1.6.0. All parameters displayed very different distributions, suggesting that the choice of the priors had very little effect on the obtained values.

### Protein family clustering and annotation

All 113 proteomes from the Legio93 set were clustered into protein families using OrthoMCL 2.0.9 ^68^. All proteins were aligned to each other with the blastp variant of DIAMOND v0.9.8.109 ^69^, with the --more-sensitive option, enabling masking of low-complexity regions (--masking 1), retrieving at most 10^5^ target sequences (-k 100000), with a E-value threshold of 10^-5^ (-e 1e-5), and a pre-calculated database size (--dbsize 98280075). In the clustering step, mcl 14-137 ^70^ was called with the inflation parameter equal to 1.5 (-I 1.5), as recommended by OrthoMCL.

Annotation of the protein families obtained by OrthoMCL was performed by searching for protein accession number in a list of reference genomes (see below). If any protein in a family was present in the first genome, the family was attributed this annotation; else, the second genome was searched for any of the accession numbers, and so on. The reference list was as follow (Genbank accession numbers between parentheses): *Legionella pneumophila* subsp. *pneumophila* strain Philadelphia 1 (AE017354), *Legionella longbeachae* NSW150 (FN650140), *Legionella oakridgensis* ATCC 33761 (CP004006), *Legionella fallonii* LLAP-10 (LN614827), *Legionella hackeliae* (LN681225), *Tatlockia micdadei* ATCC33218 (LN614830), *Coxiella burnetii* RSA 493 (AE016828), *Coxiella* endosymbiont of *Amblyomma americanum* (CP007541), *Escherichia coli* str. K-12 substr. MG1655 (U00096), *Francisella tularensis* subsp. *tularensis* (AJ749949), *Piscirickettsia salmonis* LF-89 = ATCC VR-1361 (CP011849), *Acidithiobacillus ferrivorans* SS3 (CP011849), *Neisseria meningitidis* MC58 (AE002098), *Rickettsiella grylli* (NZ_AAQJ00000000.2). *Legionella* effector orthologous groups were identified by comparing protein accession numbers with the list provided by ^20^.

### Ancestral reconstruction

The flow of genes in the order *Legionellales* was analyzed by reconstructing the genomes of the ancestors. The ancestral reconstruction was performed with Count v10.04 ^24^, which implements a maximum-likelihood phylogenetic birth- and death-model. The Bayesian input tree was obtained from the Legio93 dataset (see above) and the OrthoMCL families as described above. The rates (gain, loss, and duplication) were first optimized on the OrthoMCL protein families, to allow different rates on all branches. The family size distribution at the root was set to be Poisson. All rates and the edge length were drawn from a single Gamma category, allowing 100 rounds of optimization, with a 0.1 likelihood threshold. Reconstruction of gene flow was then performed by using posterior probabilities.

### Analysis of the Type IVB secretion system

Homologs of the Type IVB secretion system were identified with MacSyFinder v1.0.3 ^71^. We used the pT4SSi profile, corresponding to the Type IV Secretion System encoded by the IncI R64 plasmid and *Legionella pneumophila* (T4BSS; also referred to as MPF_I_) ^72^, as query to search all proteomes in the Gamma105 and the Legio93 datasets. This profile includes models for the 17 proteins of the IncI R64 plasmid that have a homolog in *L. pneumophila*. The profile does not cover 6 of the Dot/Icm proteins shared among *L. pneumophila* and *C. burnetii* (IcmFHNQSW). MacSyFinder was run in “ordered_replicon” mode, and using the following HMMer options: --coverage-profile 0.33, --i-evalue-select 0.001. A T4BSS was identified if, within 30 neighboring genes: (i) a IcmB/DotU/VirB4 homolog was present, (ii) at least 6 out of the other 16 were also present, (ii) no relaxase (MOBB) were identified. Since the Icm/Dot system is often split in several operons, not all proteins could be identified by MacSyFinder in “ordered_replicon” mode, but most were since the operon which contains IcmB/DotU/VirB4 is the largest.

To inspect the gene organization further in a selection of organisms, a subset of 12 representative genomes were aligned with tblastx 2.2.30 ^73^, with an E-value threshold of 0.01, not filtering the sequences for low complexity regions, and splitting the genomes in smaller pieces when reaching the maximum number of hits between two segments. Results were first visualized in ACT ^74^, and a final version of the comparison was drawn with genoPlotR ^75^. In the cases where MacSyFinder failed to detect a T4BSS, a similar visual inspection of the comparison of the genome with both *L. pneumophila* and *C. burnetii* was performed.

For the genomes where the T4BSS was detected by MacSyFinder, twelve proteins (IcmBCDEGKLOPT, DotAC) were retrieved, aligned with mafft-linsi v7.273 ^44^, trimmed with trimAl v1.4.rev15 ^76^, using the -automated1 setting, and concatenated. The concatenated alignment was used to infer a maximum-likelihood tree with IQ-TREE v1.6.8 ^51^, forcing the use of the LG ^52^ substitution matrix, but letting ModelFinder ^56^ select the across-site categories, which resulted in the C60 mixture model ^53^; 1000 ultrafast bootstraps were drawn ^55^. For the genomes not belonging to *Legionellales* sl (R64, *Fangia hongkongensis*, *Acidithiobacillus ferrivorans*, *Acidiferrobacter thiooxydans* and *Alteromonas stellipolaris*), only a few proteins were retrieved by the automated approach. A preliminary tree displayed very long branches, and the procedure above was repeated without these four genomes. For the 12 genomes for which the T4BSS was manually annotated, a tree based on the alignment of 26 proteins belonging to the T4BSS was performed as above, resulting in a LG+F+I+G4 model being selected.

### Data availability

The raw and assembled data for the *Aquicella* genomes is deposited at the European Nucleotide Archive (ENA) under study accession PRJEB29684. The assemblies for *A. lusitana* and *A. siphonis* have the accessions GCA_902459475 (replicons LR699114.1-LR699118.1) and GCA_902459485 (replicons LR699119.1-LR699120.1).

MAGs, genomes, associated proteomes and alignments underlying the trees presented in this contribution are available at Zenodo, with DOI 10.5281/zenodo.3543579.

## Acknowledgments

This work was supported by the Swedish Research Council [2017-03709 to L.G.], Science for Life Laboratory [SciLifeLab National Project 2015 to L.G.], and the Carl Tryggers Foundation [CTS 15:184, CTS 17:178 to L.G.].

We would like to thank Thijs Ettema and Lisa Klasson for constructive discussion; Jennah Dharamshi and Lina Juzokaite for help with the 16S amplicon protocol; Laura Eme for help with dating issues; Joran Martijn and Julian Vosseberg for the help with the Tara Ocean binning; and Jennifer Ast for her help with editing the manuscript. We would also like to thank the Ag1000G project for releasing early their data.

## Author contributions

L.G. conceived the study and drafted the manuscript. E.H. performed database screening and metagenomics; T.A. analyzed the T4BSS and effectors; E.H. and L.G. performed phylogenomics. All authors contributed to writing the final manuscript and approve it.

## Supplementary Figures

**Supplementary Figure 1:**
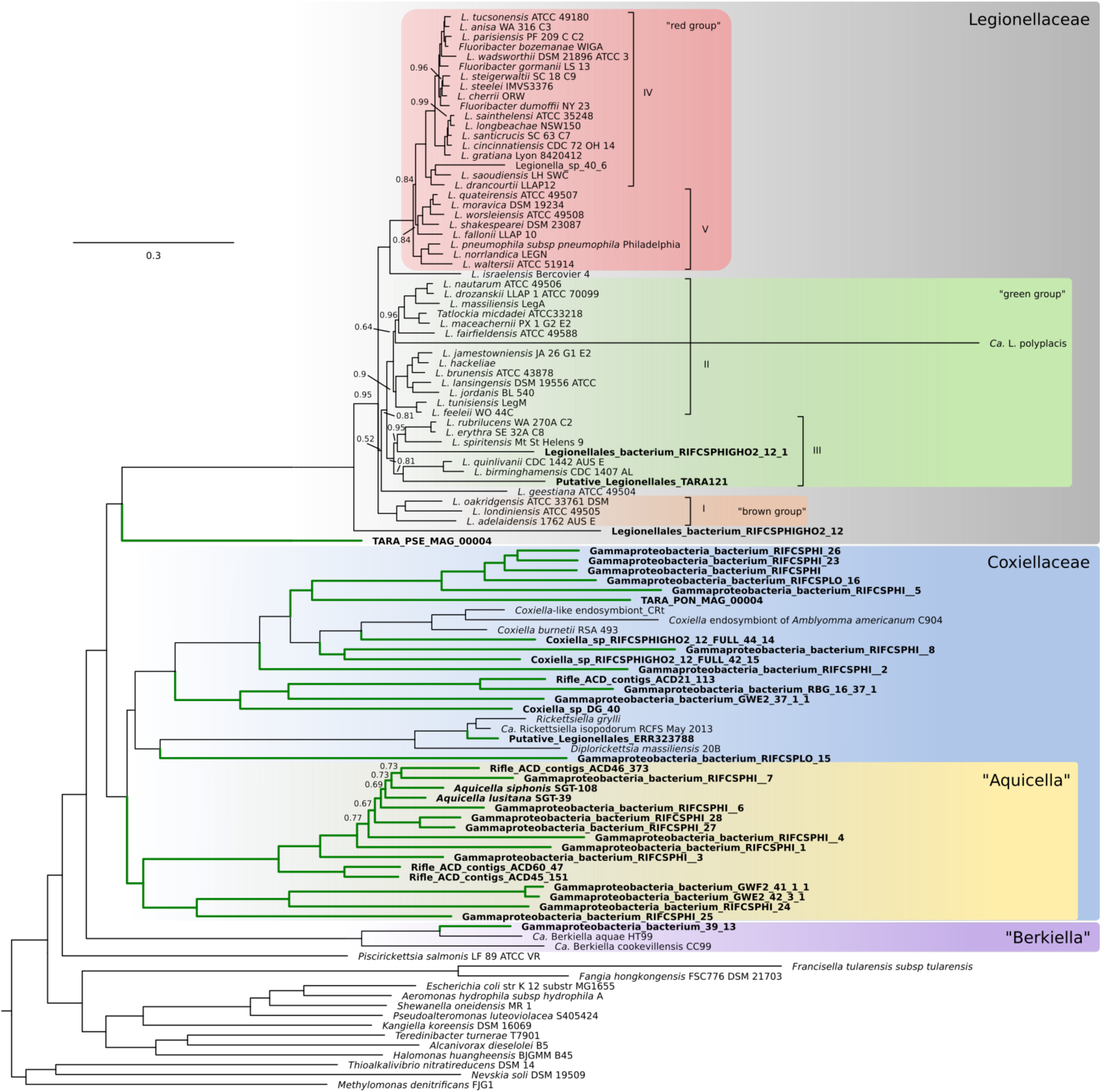
Bayesian phylogeny of the order *Legionellales*. Based on a concatenated amino-acid alignment of 105 single-copy orthologs, the tree was inferred with PhyloBayes using a GTR+CAT model. The tree encompasses 93 *Legionellales* genomes and is rooted with an outgroup of 4 genomes from *Proteobacteria* other than *Gammaproteobacteria*. Numbers on branches show posterior probabilities (pp). Branches without number indicate pp = 1. Terminal nodes in bold indicate taxa recovered in this study. Green branches indicate clades where no genomic data was available prior to this study. The scale shows the average number of substitutions per site. The colored groups and Roman numerals in the *Legionellaceae* correspond to the groupings in ^20^ and ^21^, respectively.

**Supplementary Figure 2:**
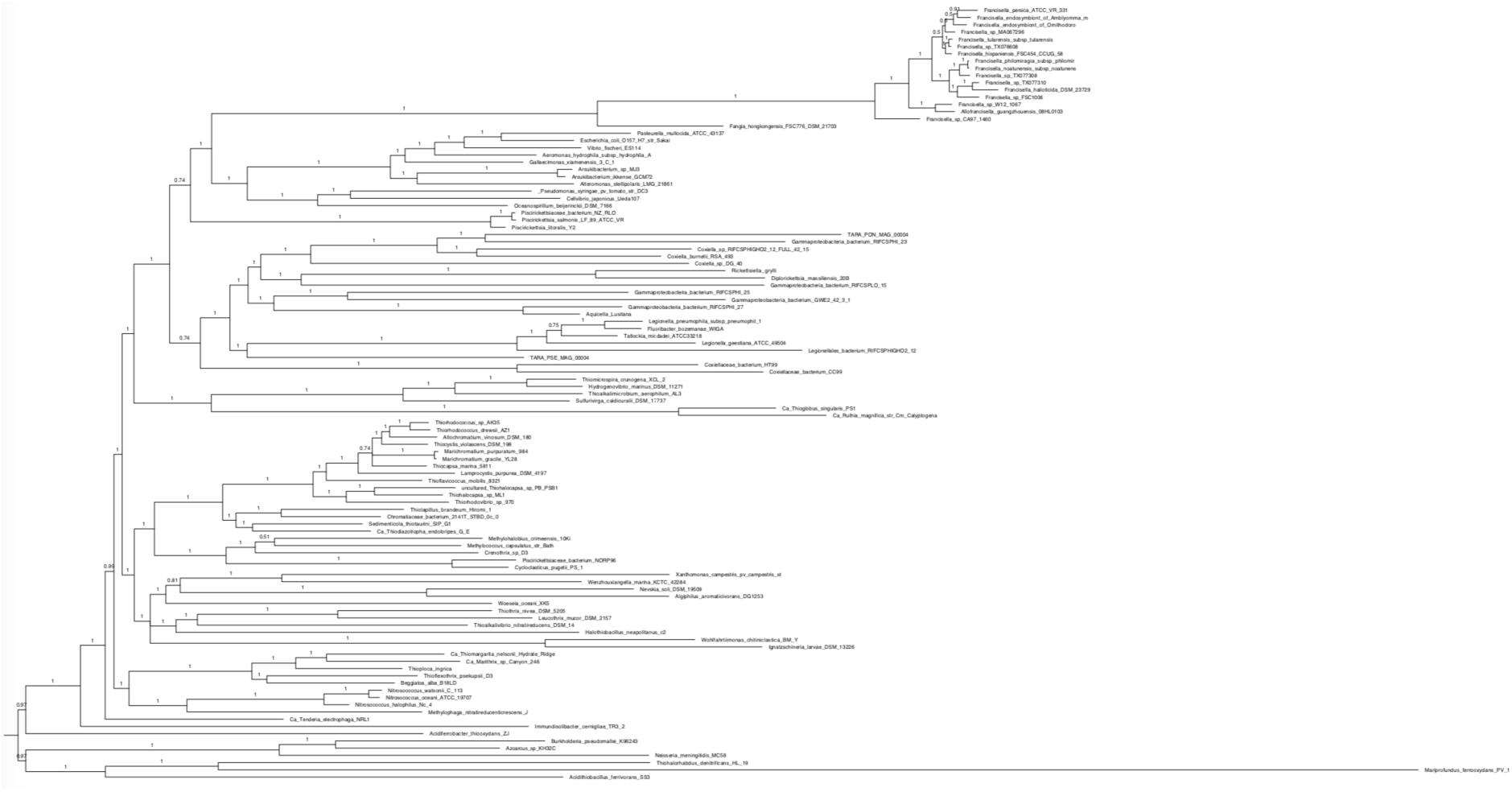
Bayesian estimation of the phylogenetic relationships of 105 *Gammaproteobacteria*. The tree is inferred from the Gamma105 dataset, which encompasses 110 genomes, including 105 *Gammaproteobacteria* and 5 other *Proteobacteria.* The selected *Gammaproteobacteria* include 22 *Legionellales*, 19 *Chromatiaceae*, 17 *Francisellaceae* and 4 *Piscirickettsiaceae*. Four phylobayes chains were run from the concatenated BMGE-trimmed alignments of 104 protein markers, using a CAT+GTR model. Numbers on the branches represent the posterior probability. The tree was rooted with the five outgroup genomes. This tree was used as an input for the time-constrained tree show on Fig. 4.

**Supplementary Figure 3:**
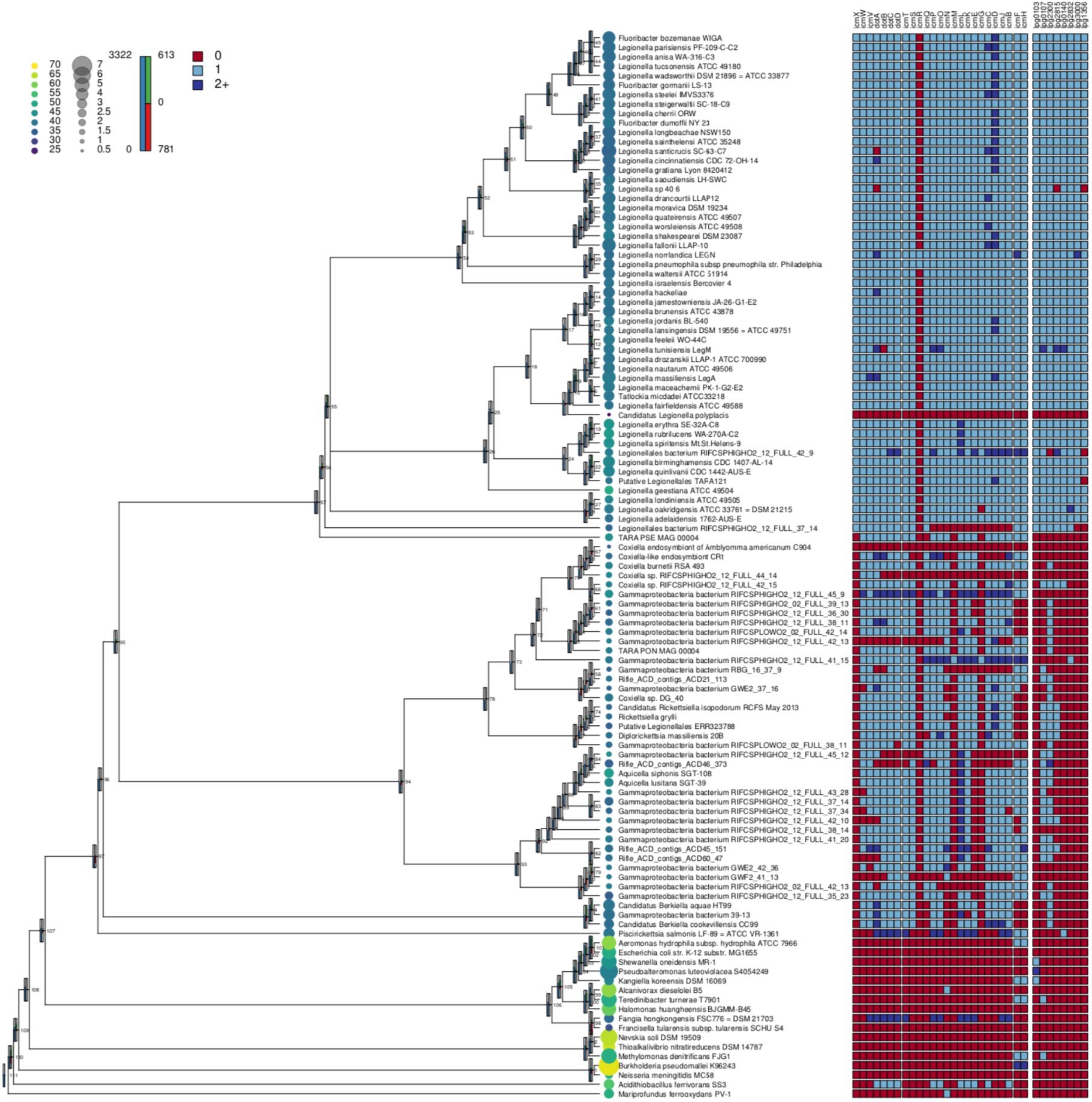
Gene flow analysis in the *Legionellales* and other *Gammaproteobacteria* genomes, with focus on the T4BSS system and the *Legionella*-conserved effectors. At the left of each node, barplots depict the number of gene families (blue, ranging from 0 to 3322) inferred in that ancestor, as well as the number of family gains (green, from 0 to 613) and losses (red, from 0 to 781) occurring on the branch leading to that ancestor. To the right of each terminal node, a circle depicts the size of the corresponding genome (proportional to the diameter of the circle) and its G+C content (color of the circle). On the right panel, squares represent proteins included in the T4BSS (separated according to the three operons present in R64 ^19^) and the eight effectors conserved in *Legionella*. The colors represent the number of homologs of each gene in the genome: red, absent; light blue, one copy; dark blue, several copies.

**Supplementary Figure 4:**
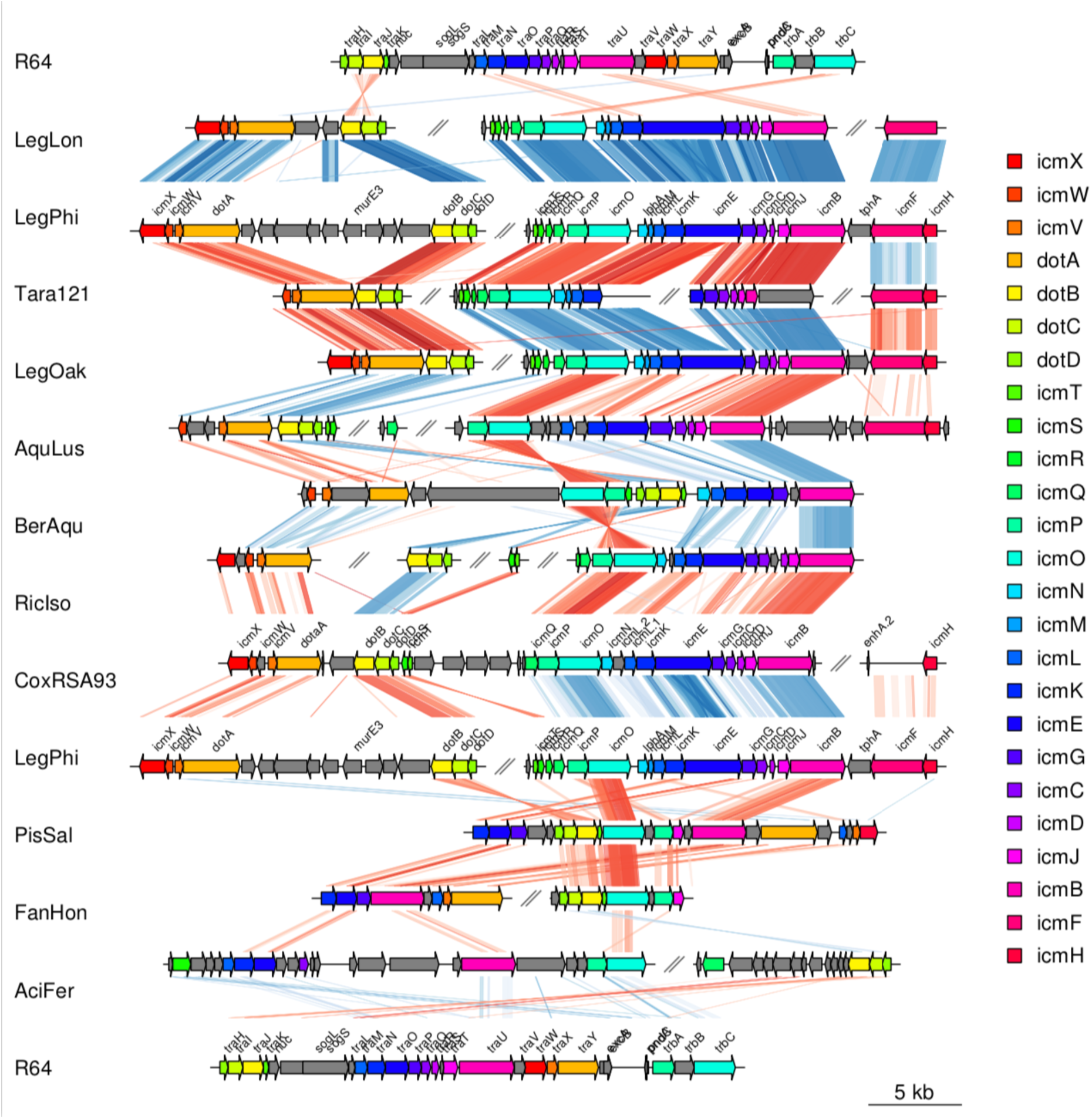
Collinearity of the T4BSS in *Legionellales*. Genes are colored as they appear in *Legionella pneumophila* Philadelphia (LegPhi). Areas between the rows display similarities between the sequences, as identified by tblastx. The sequences displayed here in addition are the IncI plasmid R64 from *Salmonella enterica* serovar Typhimurium (R64), *Legionella longbeacheae* (LegLon), a putative Legionellales TARA121 (Tara121), *Berkiella aquae* (BerAqu)*, Rickettsiella isopodorum* (RicIso), *Coxiella burnetii* RSA93 (CoxRSA93), *Piscirickettsia salmonis* (PisSal), *Fangia hongkongensis* (FanHon) and *Acidithiobacillus ferrivorans* (AciFer).

**Supplementary Figure 5:**
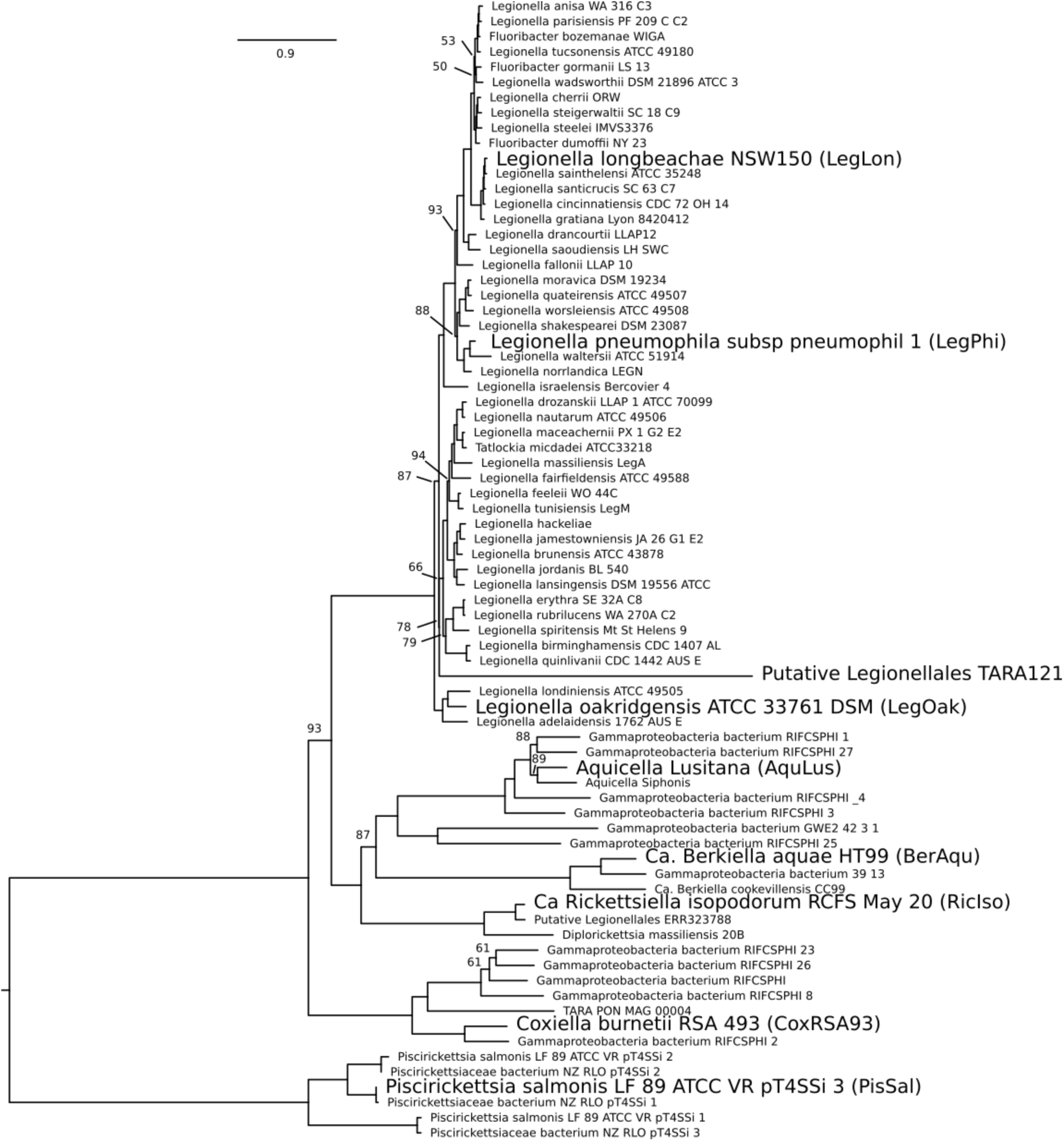
Maximum-likelihood phylogeny of the T4BSS in the *Legionellales*. The phylogeny is based on 12 genes of the T4BSS automatically detected by MacSyFinder. Number on branches represent the percent of bootstrap support. Branches without number indicate 100% bootstrap support. The scale represents the number of substitutions per site. Taxa in bold were selected for manual curation and are represented in **Supp. Figure 6** and **4**.

**Supplementary Figure 6:**
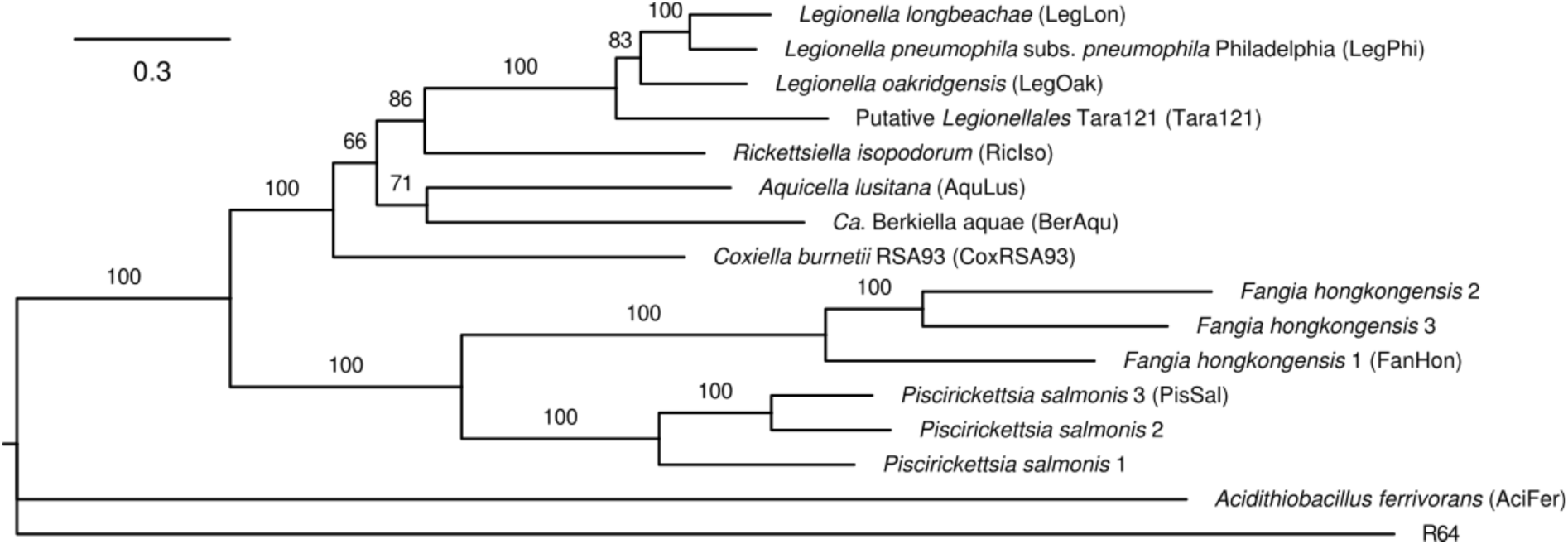
Maximum-likelihood phylogeny of the T4BSS. The phylogeny is based on the concatenation of the alignments of the 25 genes displayed on Fig. 2 and **Supp. Figure 4**, manually curated. Numbers on branches indicate the percentage of bootstrap support. The scale represents the number of substitutions per site.

**Supplementary Figure 7:**
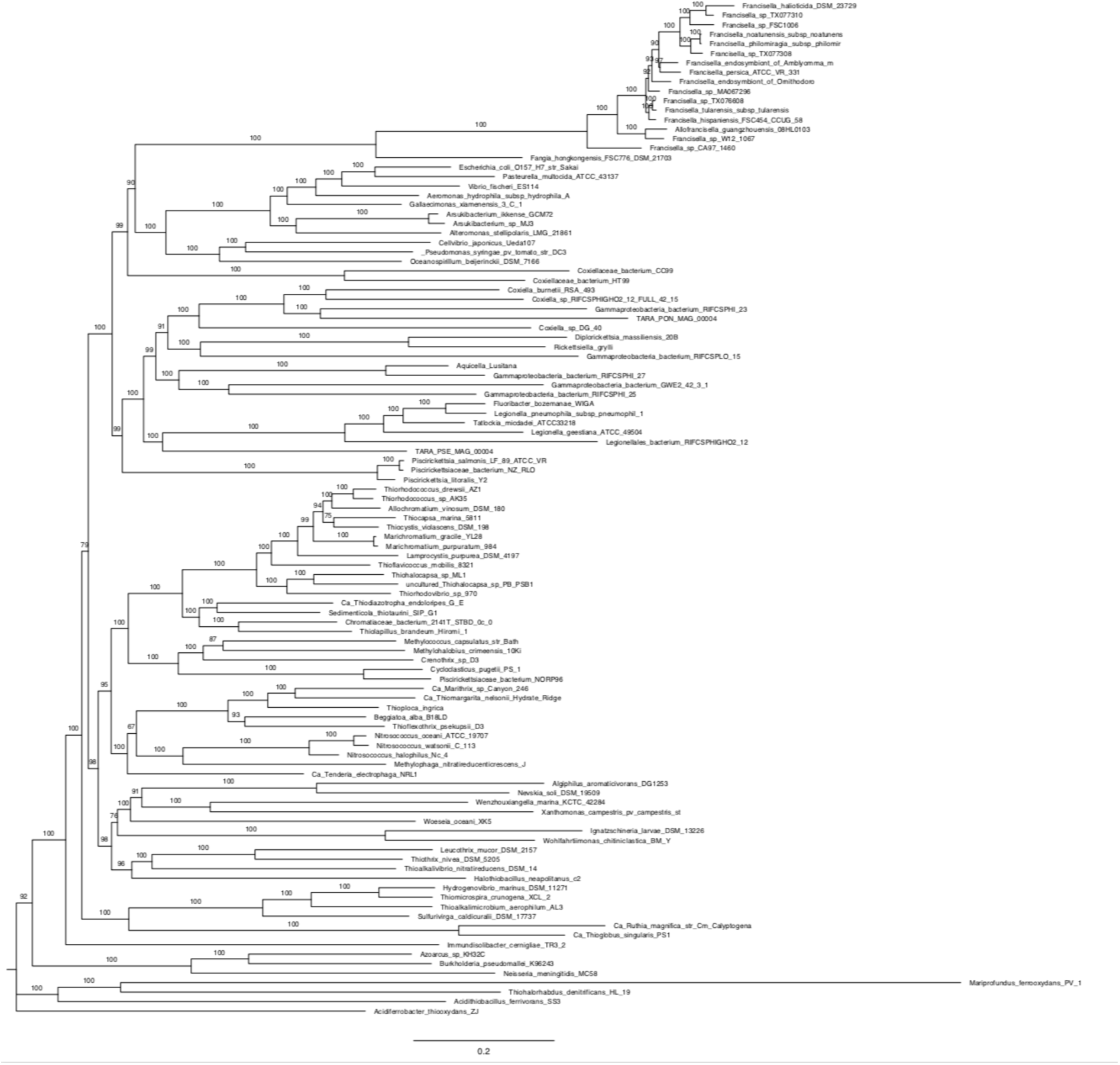
Maximum-likelihood tree of 105 *Gammaproteobacteria*. The underlying alignment is the same as in **Supp. Figure 2**. The tree was inferred with IQ-tree, using the LG substitution matrix, empirical codon frequencies, four gamma categories, the C60 mixture model and PMSF approximation. A thousand UltraFast bootstrap were drawn. The numbers on the branches represent the percentage of bootstrap tree supporting that branch. The scale represents the average number of substitutions per site.

**Supplementary Figure 8:**
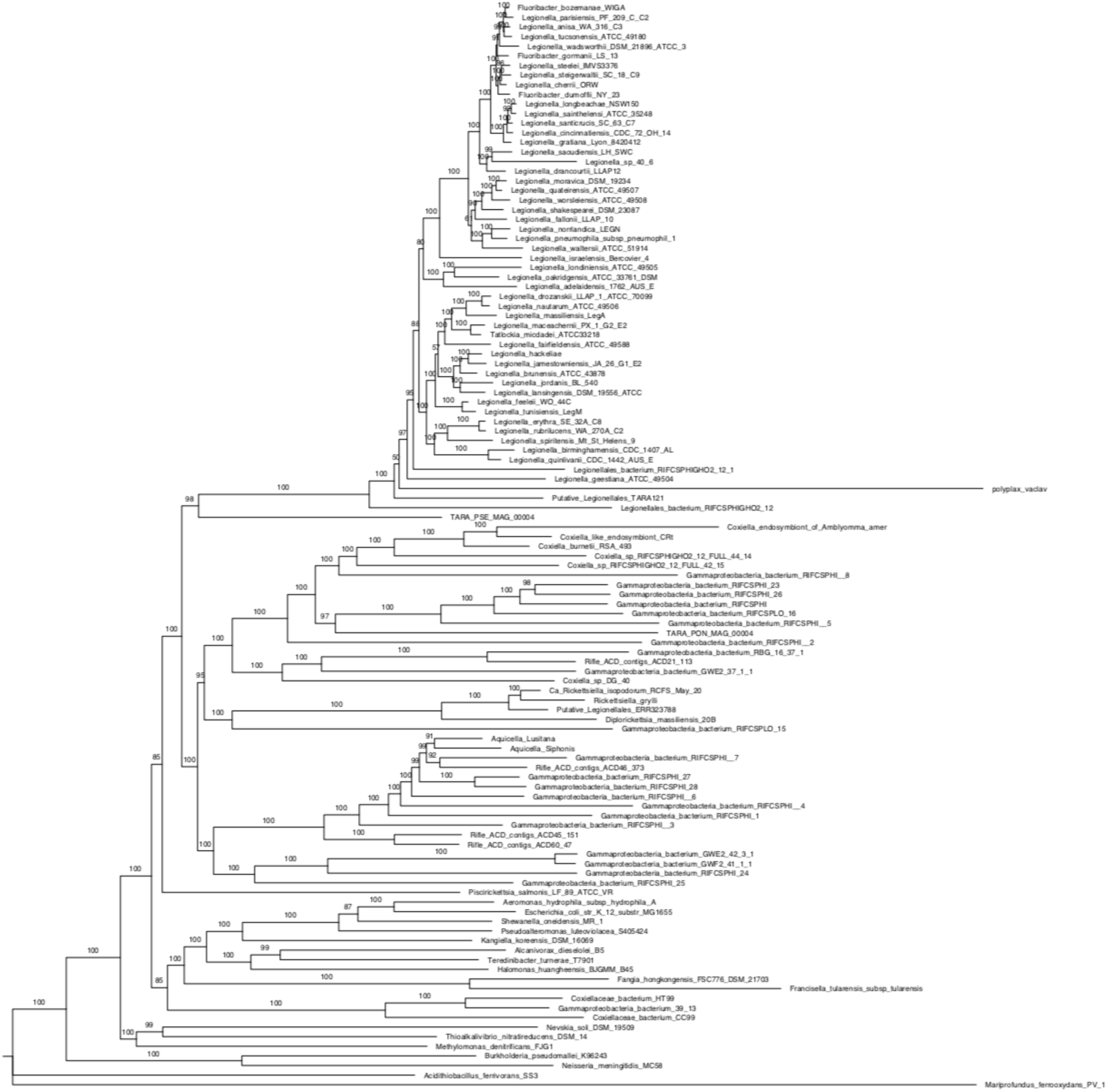
Maximum-likelihood tree of the *Legionellales*. The underlying alignment is the same as in **Supp. Figure 1**. The tree was inferred with IQ-tree, using the LG substitution matrix, empirical codon frequencies, four gamma categories, the C60 mixture model and PMSF approximation. A thousand UltraFast bootstrap were drawn. The numbers on the branches represent the percentage of bootstrap tree supporting that branch. The scale represents the average number of substitutions per site.

## Supplementary tables

**Supplementary Table 1.**
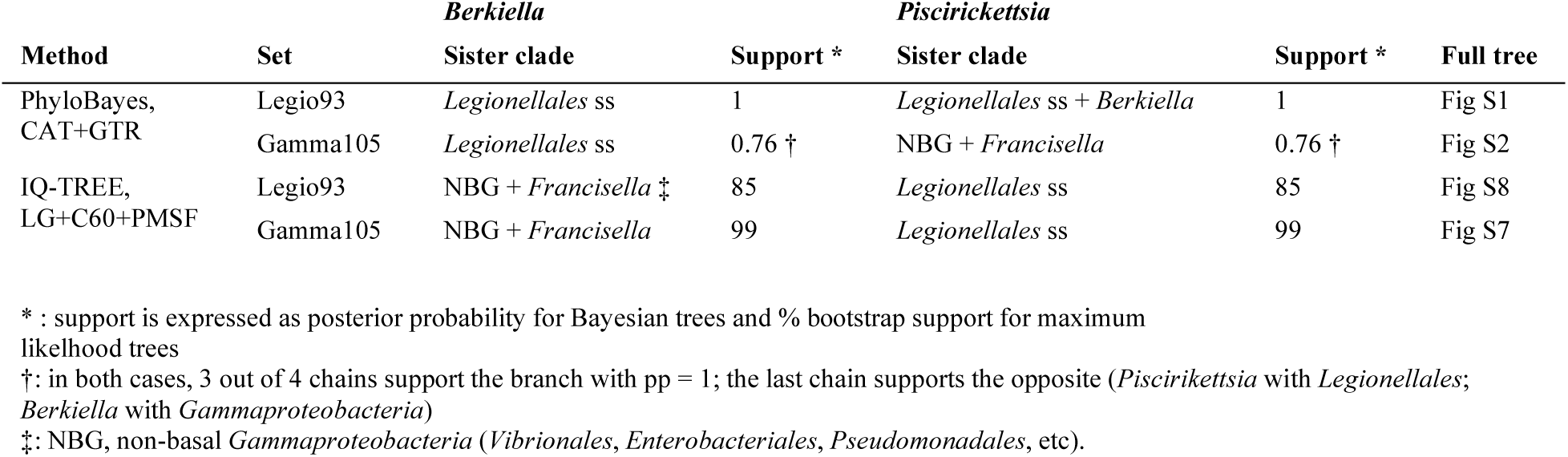
Summary of the phylogenetic placement of *Berkiella* and *Piscirickettsia* according to different methods.

| Method | Set | *Berkiella* | Support * | *Piscirickettsia* | Support * | Full tree |
| --- | --- | --- | --- | --- | --- | --- |
|  |  | Sister clade |  | Sister clade |  |  |
| PhyloBayes,<br>CAT+GTR | Legio93 | *Legionellales* ss | 1 | *Legionellales* ss + *Berkiella* | 1 | Fig S1 |
|  | Gamma105 | *Legionellales* ss | 0.76 † | NBG + *Francisella* | 0.76 † | Fig S2 |
| IQ-TREE,<br>LG+C60+PMSF | Legio93 | NBG + *Francisella* ‡ | 85 | *Legionellales* ss | 85 | Fig S8 |
|  | Gamma105 | NBG + *Francisella* | 99 | *Legionellales* ss | 99 | Fig S7 |

\* : support is expressed as posterior probability for Bayesian trees and % bootstrap support for maximum likelihood trees
†: in both cases, 3 out of 4 chains support the branch with pp = 1; the last chain supports the opposite (*Piscirickettsia* with *Legionellales*; *Berkiella* with *Gammaproteobacteria*)
‡: NBG, non-basal *Gammaproteobacteria* (*Vibrionales*, *Enterobacteriales*, *Pseudomonadales*, etc).

**Supplementary Table 2 (Separate file).**

Ancestral reconstruction of the *Legionellales*. Output from Count. For each taxon and ancestor, the table shows the estimated probabilities of families present, present in multiple copies (multi) gained, lost, expanded and contracted at each node. Ancestor IDs (numbers) are shown on the tree on **Supp. Figure 3**.

**Supplementary Table 3 (Separate file).**

Ancestral reconstruction of the genes in T4BSS and conserved effectors in *Legionellales*. For each of these genes (columns) and each of the genomes and ancestors (rows), the table shows the number of copies, the probability of presence, presence in multiple copies, gain, loss, expansion and contractions (in different blocks of rows). Ancestor IDs (numbers) are shown on the tree on **Supp. Figure 3**.

**Supplementary Table 4.**
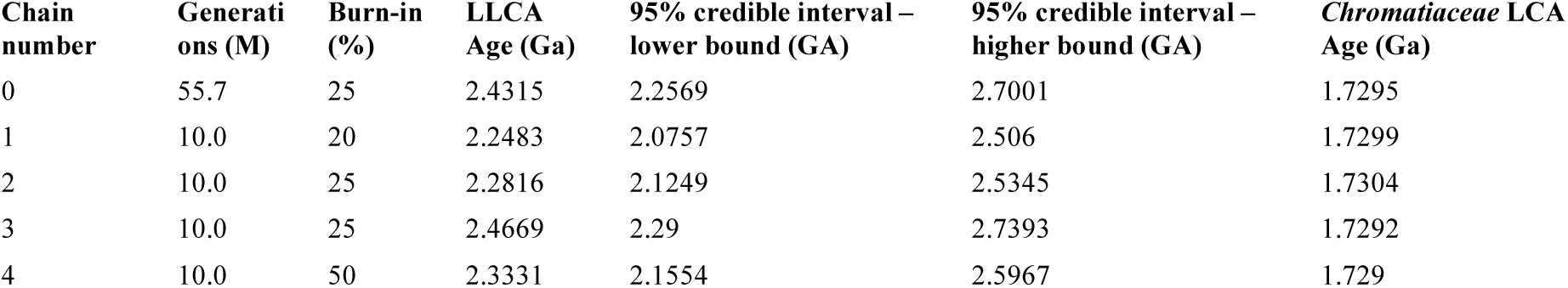
Summary of the age of LLCA in different BEAST runs.

**Supplementary Table 5:**
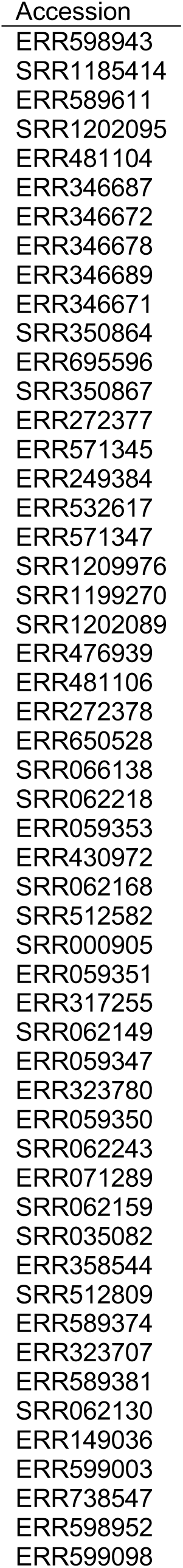
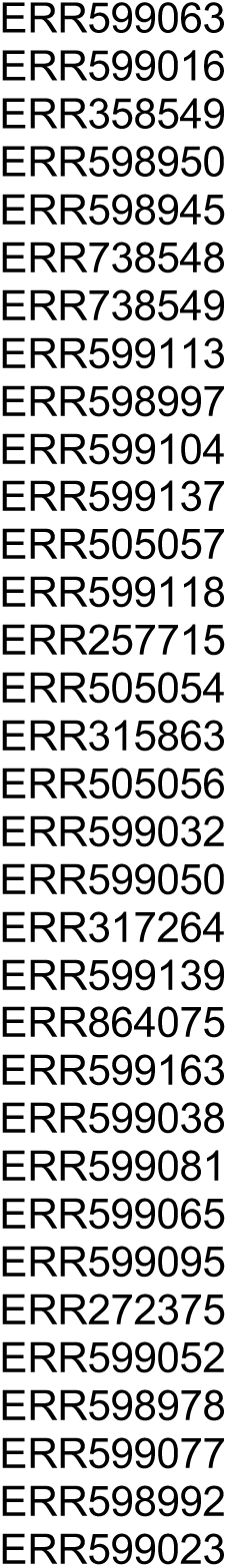
Metagenomes containing high fractions of reads mapping to *Legionellales* 16S rDNA.

**Supplementary Table 6 (Separate file).**

Gamma105 dataset, genomes used to place *Legionellales* in the *Gammaproteobacteria*.

**Supplementary Table 7 (Separate file).**

Legio93 dataset, genomes used to reconstruct the *Legionellales* genomic tree.

