## Supplementary figures and images for "Host-adaptation in *Legionellales* is 2.4 Ga, coincident with eukaryogenesis"

### Supplementary Figure 3

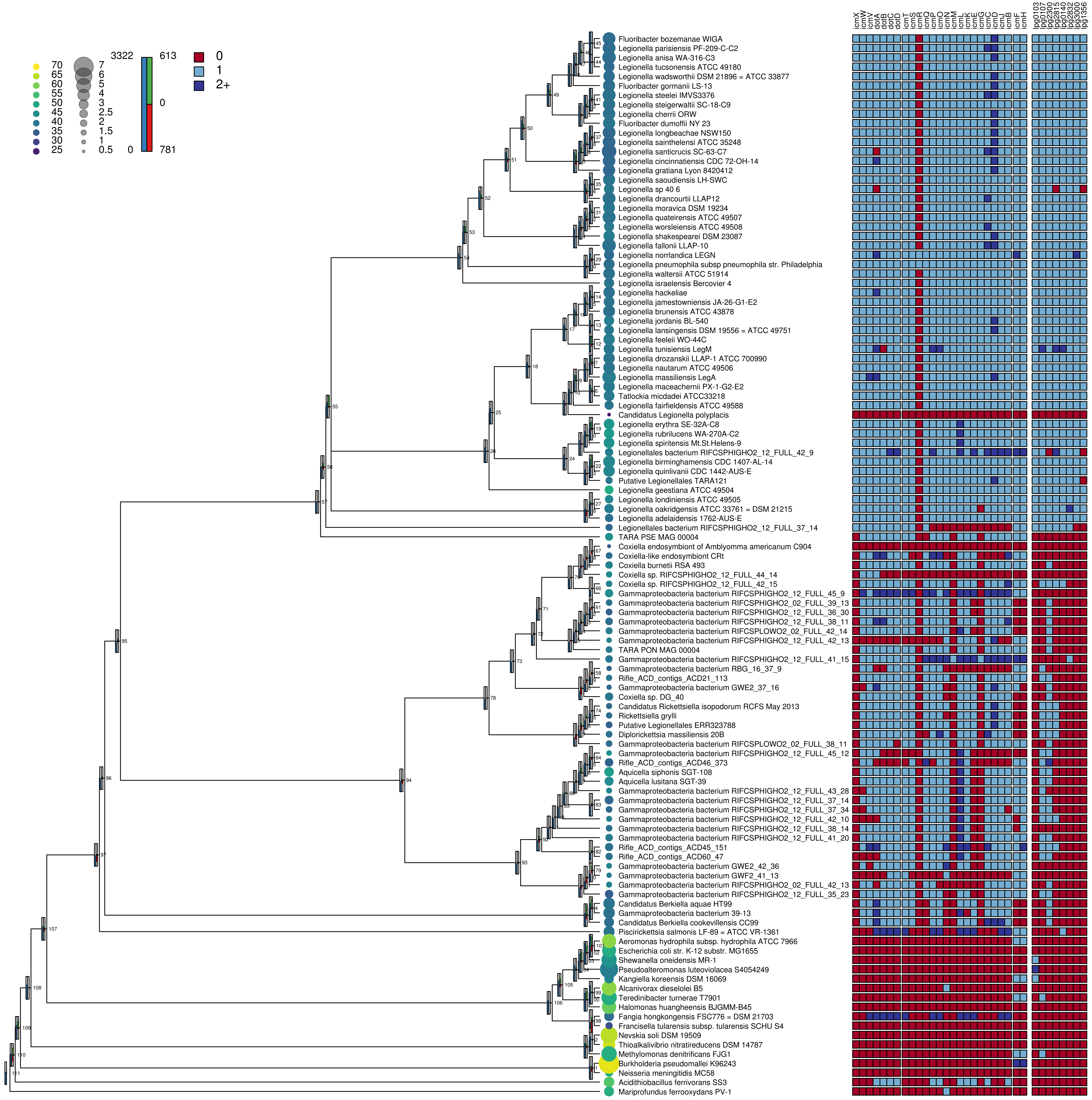
